# Identification and Quantification of Paclitaxel and its Metabolites in Human Meconium from Newborns with Gestational Chemotherapeutic Exposure

**DOI:** 10.1101/306274

**Authors:** Elyce Cardonick, Robert Broadrup, Peining Xu, Mary T. Doan, Helen Jiang, Nathaniel W. Snyder

## Abstract

**Objective:** Cancer diagnosis during pregnancy occurs in 1 out of 1000 pregnancies with common malignancies including breast and hematological cancers. Fetal exposure to currently utilized agents is poorly described. We directly assessed fetal exposure by screening meconium from 23 newborns whose mothers had undergone treatment for cancer during pregnancy.

**Study Design:** Meconium was collected from newborns whose mothers were diagnosed with cancer during pregnancy and underwent chemotherapy in the second or third trimester as part of the Cancer and Pregnancy Registry. We conducted screening of 23 meconium samples for chemotherapeutics and known metabolites of chemotherapeutics by liquid chromatography-high resolution mass spectrometry (LC-HRMS). Putative identification of paclitaxel and/or its metabolites was made in 8 screened samples. In positively screened samples, we quantified paclitaxel, 3’-p-hydroxypaclitaxel, and 6α-hydroxypaclitaxel by stable isotope dilution-LC-HRMS.

**Results:** Mean levels of paclitaxel were 399.9 pg/mg in meconium samples from newborn born to mothers that underwent chemotherapy during pregnancy. 3’-p-hydroxypaclitaxel and 6α-hydroxypaclitaxel mean levels were 105.2 and 113.4 pg/mg meconium, respectively.

**Conclusion:** Intact paclitaxel, and at least two of its major metabolites were detected in meconium, providing unambiguous confirmation of human fetal exposure. Variability in meconium levels between individuals may indicate a potential for reducing fetal exposure based on timing, dosing, and individual characteristics. This preliminary study may provide an efficient approach for examining the effects of cancer diagnosis during pregnancy on other outcomes by providing a measure of direct fetal exposure.

## Introduction

Cancer diagnosis occurs in 1 in 1000 pregnancies. The most common malignancies complicating pregnancy are breast cancer, Hodgkin’s and Non-Hodgkin’s lymphoma, and melanoma (1). The most common agents used during pregnancy include doxorubicin, cyclophosphamide, epirubicin, 5-fluouracil and more recently paclitaxel. Studies of effects of chemotherapy exposure during the second and third trimester have been reassuring, where the majority detail the physical appearance at birth and general health during the first year of life (2-6). A limited number of studies provide longer follow up with developmental and physical evaluation of children exposed *in utero* to chemotherapy up until the late teen years and young adulthood (7- 9). In all existing studies, study design has used inclusion by maternal chemotherapy during pregnancy correlated with post-natal growth and development of the children without a direct measure of fetal exposure and internalized dose to the fetus.

Pharmacokinetics of drugs in an obstetric population is complicated by changes in the mothers’ pharmacokinetic parameters and in the unique placental/fetal compartments. Differential toxicokinetics and timing of exposure impacts the potential effects from maternal exposure, placental exposure, and direct fetal exposure (10-13). Lipophilicity, affinity for transporters, molecular size of the agent, and variations in biotransformation lead to differential disposition across the maternal/fetal compartments and thus differential exposure for mother versus child. Although maternal exposure is often quantified by urine and blood analysis, direct fetal exposure is more difficult to ascertain and most would involve invasive measures such as amniocentesis or cordocentesis each of which carries risk to the ongoing pregnancy. Studies in non-human primate pregnant baboon model transplacental disposition of chemotherapeutics was possible and fetal exposure to taxanes (paclitaxel and docetaxel) occurred hours after infusion (14). Longer timepoints with docetaxel indicated a fetal reservoir, with higher fetal than maternal liver concentrations 72 hours after infusion. However, it is unknown whether fetal exposure occurs in human pregnancy and for which chemotherapeutics.

Meconium, the first stool of a newborn that passes in the first few days after birth, begins accumulation around the 13^th^ week of gestation. For this reason, it provides a unique window into gestational metabolism and exposures. Meconium also has the advantage of non-invasive collection, making it a useful biosample in epidemiologic studies of humans(15). This has been used to quantify maternal use of drugs of abuse and environmental exposures during pregnancy (15), fetal exposure to anti-retrovirals (16), metabolomics during gestational diabetes (17), and fetal sex steroid metabolism (18). To fill in the gap of the knowledge of human exposure *in utero* to chemotherapeutic agents we employed liquid chromatography-high resolution mass spectrometry (LC-HRMS) to analyze meconium sampled from newborns whose mothers underwent chemotherapy during pregnancy. Since no previous information was available on human meconium analysis for chemotherapeutics to facilitate a directly targeted approach, we used a two-step approach where we screened for the presence of major chemotherapeutics in the meconium, detected paclitaxel, and then quantified paclitaxel and its major metabolites in the positive samples.

## Materials and Methods

### Study population

This study was approved under the Cooper Health System IRB #1-028EX maternal cancer diagnosis and treatment during pregnancy: Longitudinal follow-up of child development and maternal psychological well being. Since the Cancer and Pregnancy Registry was first created, 382 women have voluntarily enrolled and provided diagnostic and treatment information regarding their cancer during and after pregnancy. Of these women, 268 have received chemotherapy during pregnancy. Since starting the current project, 30 women have enrolled and received chemotherapy, and 23 have agreed to collect meconium at the time of delivering their child. The majority of women in the study (20) were treated for breast cancer, 1 for colon cancer and 2 for Hodgkin’s Lymphoma. Mean gestational age at the first chemotherapy treatment was 20 +/- 4.8 weeks, and mean number of days between last treatment during pregnancy and day of birth was 36 +/- 20.6 days.

### Biosample Collection

Patients treated with chemotherapy during pregnancy who consented to participate in this study were instructed to collect a single sample from the first 3 stools on day one of life. Meconium was collected on day 1 of life in all cases. Collection was performed by the mother herself, sent frozen in provided pre-labeled packaging and when received, were stored at −80°C until batched and shipped to the laboratory for analysis. Meconium was also collected from healthy newborns from the Hospital of the University of Pennsylvania in collaboration with the March of Dimes prematurity research center. Samples were collected by nurses in the hospital, stored at 4°C for under 24 hours, and then at −80°C until use. These samples were used for analytical method development, as negative control meconium, and as a matrix for spiking in analytes to generate quality control samples.

### Chemicals

Water, methanol, ethyl acetate, methyl tert-butyl ether (MTBE), acetonitrile, and acetic acid were Optima LC-MS grade solvents from Fisher Scientific (Pittsburgh, PA). Sodium chloride and paclitaxel were from Sigma-Aldrich (St. Louis, MO) and ^13^C_6_-paclitaxel, 3’-p-hydroxypaclitaxel and 6α-hydroxypaclitaxel were from Toronto Research Chemicals (Toronto, Canada).

## Meconium Analysis

Analysts were blinded to all sample identities during processing, including untargeted and targeted analysis. 3 aliquots of meconium were analyzed per newborn. Meconium analysis was conducted by liquid-liquid extraction with either ethyl acetate or MTBE. Briefly, 50 mg (average 50.6 mg) of meconium was weighed into 10 mL screw cap glass tubes. Wet weight was recorded to 0.1 mg on a balance with tolerance to +/- 0.01 mg. Samples were spiked with an internal standard to adjust for variation in extraction and analysis. For screening, 500 pg diclofenac (50 µL of 10 pg/ µL stock in methanol) was used as an internal standard to adjust for consistency of extraction and system suitability. In quantitative analysis, 500 pg of ^13^C_6_-paclitaxel (50 µL of 10 pg/ µL stock in methanol) was used as the internal standard. During screening experiments, ethylacetate and MTBE extraction were tested since a variety of extractions have been reported from other matricies for the chemotherapeutics studied here. Briefly, ethylacetate extraction was conducted by adding 100 µL of saturated NaCl solution in water, 900 µL of water, sonicating in a bath sonicator for 10 minutes then adding 8 mL of ethyl acetate and vortexting for 30 minutes. MTBE extraction was similar except that 8 mL of MTBE was used. The organic layer was then removed and evaporated to dryness under nitrogen gas. Dried residue was re-suspended in 100 µL of 80:20 water: methanol.

LC-HRMS analysis was conducted on an Ultimate 3000 quaternary UPLC equipped with a refrigerated autosampler (6° C) and a column heater (60° C) coupled to a Thermo QExactive Plus. LC separations were conducted on a Waters XBridge C18 (3.5 µm, 100 Å, 2.1 × 100 mm). A multi-step gradient at 0.3 mL/min flow with solvent A (water 1% acetic acid) and solvent B (acetonitrile 1% acetic acid) was as follows: 10% B from 0 to 1 minute, increasing to 100% B from 1 to 5 minutes, holding 100% B until 12 minutes, then the column was returned to starting conditions and re-equilibrated at 20% B from 12 to 15 minutes. This gradient was sufficient to resolve an interfering peak (retention time of 5.5 min) in meconium samples that interfered with the detection of paclitaxel by HRMS. Column effluent was diverted to waste from 0 to 0.5 minutes and from 12.5 to 15 minutes. The mass spectrometer was operated in positive ion mode with a second-generation heated electrospray ionization source (HESI-II) alternating between full scan (200-900 m/z) at a resolution of 70,000 and data independent analysis (MS/MS) at 17,500 resolution with a precursor isolation window of 0.7 m/z. In the targeted experiments MS/MS was conducted on the M+H of paclitaxel, ^13^C_6_-paclitaxel, 3’-p-hydroxypaclitaxel and 6α-hydroxypaclitaxel. Source parameters were as follows: capillary temperature, 425 °C, probe temperature, 425 °C, sheath gas, 45 arb units, aux gas, 15 arbitrary units, sweep gas, 2 arbitrary units, spray voltage 4.0 kV, S-lens, 50. Targeted data analysis was performed in Xcalibur and TraceFinder software (Thermo Fisher, San Jose, CA).

## Statistical analysis

Data was analyzed and summary statistics were calculated in Tracefinder, Excel 2016 (Microsoft), and/or with Graph Pad Prism (v7).

## Results

### Putative Detection of Paclitaxel and its Metabolites in Human Meconium by LC-HRMS

Samples from 23 newborns born to mothers who underwent chemotherapy during pregnancy were screened by LC-HRMS for the presence of chemotherapeutics and their major metabolites. Meconium from 10 healthy newborns was used for assay development; as a negative control to establish blank levels, and as a matrix to spike known quantities of analyte to serve as positive controls for calibration curves and quality controls. A chromatographic peak putatively corresponding to intact paclitaxel was detected in meconium from 7/23 meconium samples, with putative detection of two paclitaxel metabolites but no parent drug in one other sample retrospectively found after unblinding. Detection frequency was markedly lower with MTBE extraction (4/23) from meconium despite this being a common liquid-liquid extraction solvent for quantification of paclitaxel in other matricies(19).

### Quantification of Paclitaxel and its Metabolites in Positively Screened Meconium Samples

To confirm the screening results by LC-HRMS we utilized targeted stable isotope dilution-LC-HRMS using commercially available ^13^C_6_-paclitaxel instead of the surrogate internal standard diclofenac used in screening. Human metabolism of paclitaxel occurs via cytochrome P450s to hydroxylated and epoxide metabolites, with high levels of excretion in the bile as well as urinary excretion (20). Major metabolites include 3’-p-hydroxypaclitaxel, and 6α-hydroxypaclitaxel produced by CYP3A4 and CYP2C8, respectively, thus we added these analytes to our targeted assay (21). 3’-p-hydroxypaclitaxel and 6α-hydroxypaclitaxel were well resolved on our LC-HRMS method and were quantified using ^13^C_6_-paclitaxel as an internal standard (**Fig. 1A**). Standard curves over the range of 4-4,000 pg/mg meconium were linear with R^2^ values of 0.9949, 0.9964 and 0.9971 for paclitaxel, 3’-p-hydroxypaclitaxel and 6α-hydroxypaclitaxel, respectively. All positively detected samples were interpolated from these curves. Assay performance was confirmed by inter- and intra-day accuracy and precision with bias and % coefficient of variation less than 20% across quality controls of spiked meconium samples at 100, 200 and 500 pg/mg meconium (**Table 1**).

**Table 1.**
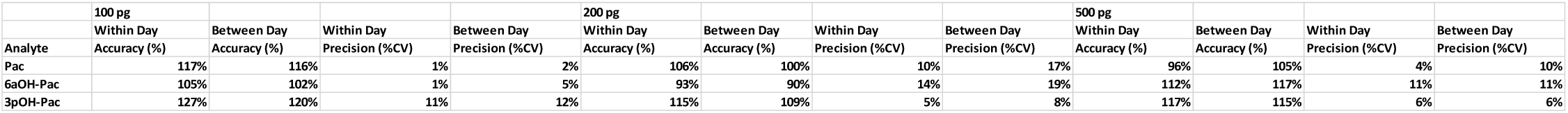
LC-HRMS assay accuracy and precision within and between days.

**Figure 1.**
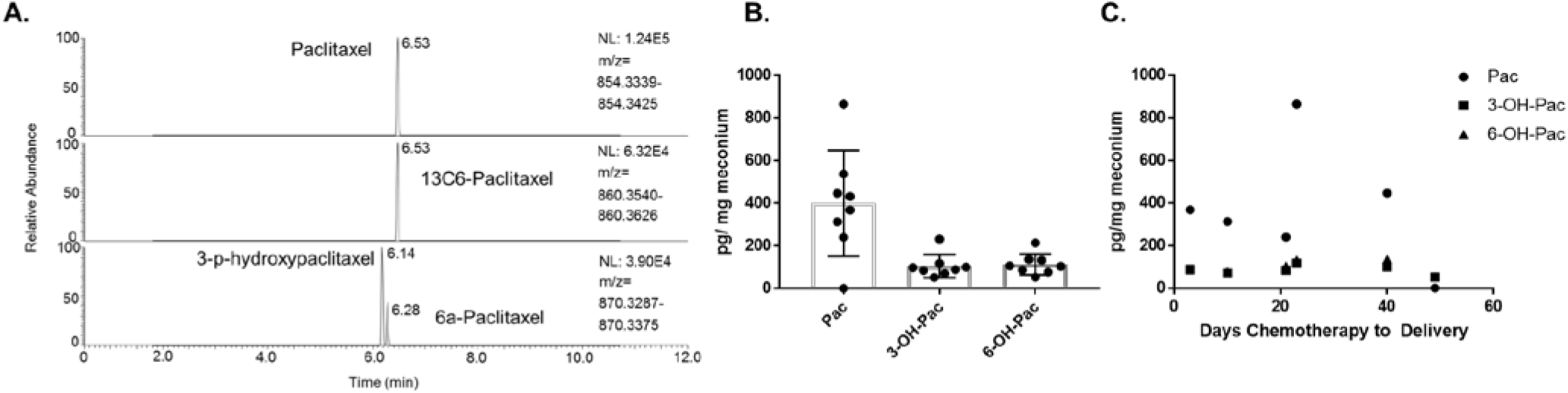
Quantification of paclitaxel in meconium. (A) Representative chromatogram of paclitaxel and its metabolites by isotope dilution-LC-HRMS in meconium. (B) Pg per mg wet weight of meconium for paclitaxel (Pac), 3’-p-hydroxypaclitaxel (3-OH-Pac), and 6α-hydroxypaclitaxel (6-OH-Pac) in 8 meconium samples. The mean of three averaged replicates per individual is shown with the standard deviation. (C) Pg per mg wet weight of meconium plotted by days of chemotherapy received before delivery.

Quantification of paclitaxel across samples, taking an average of all three aliquots, revealed a mean (standard deviation) of 399.9 (248.6) pg/mg in positively screened samples (**Fig. 1B**). 3’-p-hydroxypaclitaxel and 6α-hydroxypaclitaxel were quantified at 105.2 (54.6) pg/mg and 113.5 (48.9) pg/mg, respectively. Spearman rank correlation was used to analyze the correlations of the averaged values of paclitaxel and its metabolites in the meconium samples corresponding to the same individual. Correlation of paclitaxel with its metabolites was strong (Spearman r (p-value) for each pair paclitaxel/3’-p-OH-Pac 0.952 (p=0.00114) paclitaxel/6α-OH-Pac 0.857 (p=0.001071)) and correlation between the two metabolites was strong at 0.952 (p=0.00114).

For 6 samples where data was available on treatment schedule, levels of paclitaxel and its metabolites was plotted by days of chemotherapy received until delivery (**Fig. 1C**). Correlation between meconium levels and days of chemotherapy before delivery was negative, weak, but not statistically significant (Pearson r (95% CI bounds)= −0.2918 (−0.8921 to 0.681), −0.2507 (−0.8827 to 0.7041), −0.01241 (−0.8158 to 0.8073) for paclitaxel, 3’-p-hydroxypaclitaxel and 6α-hydroxypaclitaxel, respectively.

## Discussion

To our knowledge, this is the first measurement of chemotherapeutics in human meconium. As a preliminary study, there are severe limitations to our findings. First, we were unable to obtain any other maternal or fetal biospecimens to examine circulating maternal chemotherapeutic concentrations. Combined with a lack of dosage information and a complete accounting of covariates (other pharmaceuticals given, comorbidities, body weight, etc.) this prevents informed pharmacokinetic analysis within this study design. Second, the use of analytical screening before targeted quantification may limit the ability to detect other chemotherapeutics in meconium. However, it takes significant resources to develop and then validate methods in new biological matrices, especially complex ones such as meconium. Future studies could take advantage of the findings here to further this work with more complete data and biosample collection. The benefit of meconium, in terms of ease of sample collection with a low burden on participants and researchers, may make this feasible.

The quantification of paclitaxel and its metabolites in meconium agrees with fetal disposition quantified in previous pregnant baboon studies (14). Similarly, *ex vivo* studies of human placental transport of taxanes partially agrees with our results. Of particular relevance to interpretation in this study, Berveiller, *et al*., performed transplacental studies of paclitaxel and docetaxel in term placentas using a human perfused cotyledon placental model (13). Targeted maternal concentrations mimicked an infusion of 90 mg/m^2^ of paclitaxel and 100 mg/m^2^ of docetaxel used in clinical practice. Fetal transfer rate and placental uptake ratio were calculated as transport parameters. After perfusion, the cotyledons underwent extraction to quantify docetaxel and paclitaxel using reverse-phase high-performance liquid chromatography. Results showed that the transplacental transfer of paclitaxel and docetaxel were low and similar. Large heterogeneity was also noted across placentas, similar to the meconium variability in this current study. An explanation offered by the authors was the differential expression of p-glycoprotein (P-gp), a drug transporter which should decrease tissue accumulation in the fetus. Authors found both variability in expression according to both gestational age and between patients of similar gestational age. According to Hemauer et al, polymorphisms in the MDR1 gene on the expression affect the activity of placental P-gp and therefore P-gp-mediated active transport of paclitaxel. When authors examined 200 human placentas there was inter-individual variability and no correlation between P-gp protein expression and transport activity of paclitaxel. Consequently, investigators hypothesized that genetic variation could contribute to discrepancies between observed protein expression and activity of placental P-gp there was a gene-dose effect, whereby heterozygous placentas (wild type/variant) had intermediate level of P-gp uptake activity compared to placentas with the wild type (lowest uptake) and homozygous variants (highest uptake).(22) In this current study we did not perform genetic studies of mothers’ or children’s MDR1 gene expression.

Heterogeneity in the amount of paclitaxel and its metabolites in different individuals may indicate either differential maternal pharmacokinetics or different fetal pharmacokinetics. Variations in pharmacokinetic parameters of paclitaxel have been reported from ABCB1 polymorphism and P-gp activity (23), and variations in toxicity have been reported by CYP3A4 genotype (24). When administered intravenously over 1 to 96 hours to adults, paclitaxel has demonstrated the following pharmacokinetic characteristics: extensive tissue distribution; approximately 90 to 95% plasma protein binding, average systemic clearance varying between 87 to 503 ml/min/m^2^ (5.2 to 30.2 L/h/m^2^); and <10% renal excretion of parent drug. In recent trials in children and adults, paclitaxel elimination appeared to be saturable, the clinical importance of which would be greatest when large dosages are administered and/or the drug is infused over a shorter period. In these situations, achievable plasma concentrations are likely to exceed the constant for elimination. Thus, small changes in dosage or infusion duration may result in disproportionately large alterations in paclitaxel systemic exposure (10). Our findings also agree with the descriptions of fecal excretion of paclitaxel metabolites in adults acknowledging the major caveat that fecal elimination in adults is not equivalent to meconium deposition.

Regardless, this study should be taken into context within the clinical evidence base of the effectiveness of taxol-based chemotherapy. No data in this study suggests or assumes toxic levels of chemotherapeutics are achieved in the fetal compartment. In fact, the use of paclitaxel and docetaxel have been reported in human pregnancies with reassuring neonatal outcomes (25-32). When looking at the developmental outcomes of infants exposed to chemotherapy, prematurity had a stronger negative influence on development than chemotherapy (8). This study does suggest that meconium analysis has utility as a measure of fetal exposure in precision medicine for designing and testing dosing regimens to improve maternal delivery of chemotherapeutics. Meconium analysis may also help understand human variability in maternal pharmacokinetics and provide more statistically efficient observation of developmental outcomes after cancer during pregnancy.

